# Sparse network-based regularization for the analysis of patientomics high-dimensional survival data

**DOI:** 10.1101/403402

**Authors:** André Veríssimo, Eunice Carrasquinha, Marta B. Lopes, Arlindo L. Oliveira, Marie-France Sagot, Susana Vinga

## Abstract

Data availability by modern sequencing technologies represents a major challenge in oncological survival analysis, as the increasing amount of molecular data hampers the generation of models that are both accurate and interpretable. To tackle this problem, this work evaluates the introduction of graph centrality measures in classical sparse survival models such as the elastic net.

We explore the use of network information as part of the regularization applied to the inverse problem, obtained both by external knowledge on the features evaluated and the data themselves. A sparse solution is obtained either promoting features that are isolated from the network or, alternatively, hubs, *i.e.*, features that are highly connected within the network.

We show that introducing the degree information of the features when inferring survival models consistently improves the model predictive performance in breast invasive carcinoma (BRCA) transcriptomic TCGA data while enhancing model interpretability. Preliminary clinical validation is performed using the Cancer Hallmarks Analytics Tool API and the String database.

These case studies are included in the recently released glmSparseNet R package^1^, a flexible tool to explore the potential of sparse network-based regularizers in generalized linear models for the analysis of omics data.

## 1 Introduction

Big data have become one of the hallmarks of many areas of research (Yin and Kaynak 2015), as the associated high dimensionality of the information makes it very demanding for state-of-the-art models to extract relevant knowledge (Alyass, Turcotte, and Meyre 2015). In addition, it becomes difficult to trace which are the most significant features in the models. Although interpretation can be discarded in some areas, in a clinical setting it becomes unrealistic to use black box models to make decisions without the practitioner understanding which factors drive a decision, regardless of the accuracy of the method.

Precision medicine is characterized by combining the advances in biomedical reasearch with the availability of genomic and transcriptomic data from patients to create individualized and targeted therapies (He et al. 2013). This is now becoming possible, as the cost of sequencing is declining (Phillips, Pletcher, and Ladabaum 2015) with tumor sequencing becoming a standard procedure to plan cancer treatment (Taber, Dickinson, and Wilson 2014). The challenge is to integrate physiological, laboratory tests and molecular data in the decision making process, to better adjust the treatment course and its overall prognostic.

As the amount of information available rapidly increases in dimensionality and complexity, black box methods (e.g. neural networks, support vector machines, genetic programming) or even linear models may remain accurate but interpretation of the contribution of the most important features becomes impracticable, particularly in high-dimension settings.

A strategy for minimizing this problem is to promote a sparse solution that minimizes the number of contributing features, either by feature selection or directly in the model. The least absolute shrinkage and selection operator (Lasso) approach (Tibshirani 1996) of penalizing the *L*1-norm attempts to promote these sparse solutions. However, it is a mathematical method which has been proven to arbitrarily pick single features from subsets of correlated ones, leaving the others out of the solution. In certain conditions, there could be multiple optimal solutions for the problem, which worsens the interpretability problem. This is true not only for Lasso, but also for other methods.

Another way of constraining the solutions in order to increase model interpretability is the incorporation of prior network information in the penalization term (Ozturk et al. 2018). Recent work on network-based regularization in Cox regression, DegreeCox (Veríssimo et al. 2016), has shown promise in improving survival prediction against Lasso in ovarian cancer patients based on gene expression data, while promoting interpretability of the models obtained. DegreeCox enables the incorporation of node information by favouring particularly relevant genes, *e.g.* extracted from available gene co-expression networks or correlation/covariance networks extracted from the data themselves.

This work evaluates different network-based strategies for the generation of survival models for the breast invasive carcinoma (BRCA) transcriptomic data available from The Cancer Genome Atlas (TCGA) Data Portal. Two strategies are evaluated for the construction of the network-based penalty term: i) data-based weights attributed to each node/gene (correlation, covariance and Bayesian networks extracted from the data), with lower penalties being attributed either to hub genes, *i.e.* highly connected genes, or weakly connected or isolated genes, herein called *orphan* genes; ii) gene weights derived from external information on a protein-protein interaction (PPI) network. The increased model accuracy and interpretability obtained are expected to provide valuable insights on the biology of the diseases and point to new directions towards appropriate personalized treatment.

## 2 Methods

The proposed method to improve survival analysis by using network information is based on the Cox proportional model (Cox 1972) and the use of the degree centrality metric for graphs in order to explore the relation between the degree of a node and its weight in the solution.

We first start by describing the Cox proportional hazard model with regularization, we then describe the networks used in this study, and the way the degree was used to augment the models.

### 2.1 Cox proportional hazard model

The Cox proportional hazard model was extended (Friedman, Hastie, and Tibshirani 2010) to introduce a penalization term to the objective function using a Tikhonov (Tikhonov and Arsenin 1977) formulation. The penalization function uses *L*1 and *L*2-norms to allow for shrinkage.

Given *D* = ((***X***_1_*, Y*_1_*, δ*_1_)*, …,* (***X***_*n*_*, Y*_*n*_*, δ*_*n*_)) for *n* individuals and *p* features, where *X*_*ij*_ is the feature *j* value for individual *i*. The response variable, *Y*_*i*_, indicates the survival time for patient *i* and *δ*_*i*_ is an indicator of whether the event was observed for patient *i* (*δ*_*i*_ = 1) or not (*δ*_*i*_ = 0). The hazard function for a patient given his expression profile is given by the equation below at time *t* for subject *i* with explanatory variables ***X***_*i*_ and ***β*** = (*β*_1_*, …, β*_*p*_) the unknown parameters:

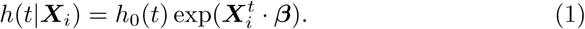

Let *t*_1_*, …, t*_*n*_ be the follow-up times for the *n* individuals. By denoting the ordered follow-up times by *t*_(1)_ *< … < t*_(*n*)_, the partial likelihood is given by the following expression

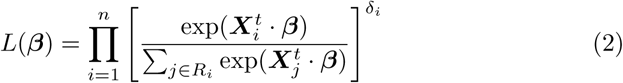

where *R*_*i*_ = *R*(*t*_(*i*)_) = {*j* : *t*_*j*_ *≥ t*_(*i*)_} denotes the risk set of all individuals that are at risk at *t*_(*i*)_, i.e., with a follow-up time greater that or equal to *t*_(*i*)_.

The negative partial log-likelihood function that can be minimized over ***β*** with the regularization parameter and function, *λ* and **Ψ**(***β***) respectively, is equal to:

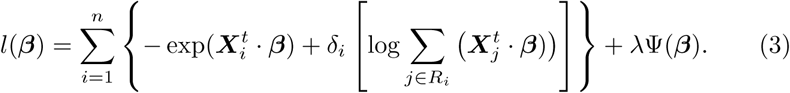

The addition of **Ψ**(***β***) to the partial log-likelihood will penalize the model with a structure that can impose a structural restriction that is data-independent.

Our goal is use **Ψ** to explore how network information can be used to individually restrict or promote the selection of a feature. We propose to use graph-based centrality measure for this purpose, using a network that models the relationships between the features, which can be generated from the data itself or use an external network described in the literature.

In this work we propose to extend DegreeCox (Veríssimo et al. 2016) by introducing sparsity in the definition of **Ψ** by adding a *L*1-norm term, also known as elastic net (Zou and Hastie 2005). The **Ψ** function is then defined as:

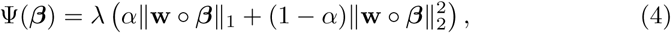

where º is the Hadamard product (or element-wise product) between the two vectors and **w** represents the degree centrality information to be included.

### 2.2 Biological networks

The biological networks explored in this work are represented as undirected graphs *G* := (*V, E*), with *V* denoting the set of nodes, and *E* the set of edges that connect pairs of nodes. In the present context of gene networks, *G* represents a relationship between expressed genes where the nodes are *p* features, with *P* := | *V* |, and edges represent a weighted relation between two genes. The graph *G* may also be represented by a *p× p* positively weighted adjacency matrix that we denote by **W**.

The topological centrality of a node is characterized by the position and relationship within the network (Freeman 1978), which can use the local information, such as neighborhood of a node, or its importance by measuring the position in the flow of information. Among the proposed methods to classify the centrality of a node are degree, betweenness and closeness centrality (Bavelas 1950; Leavitt 1951; Smith 1950).

The degree of a node is defined as the sum of adjacent weighted or un-weighted edges (Fig. 1), given by

**Figure 1:**
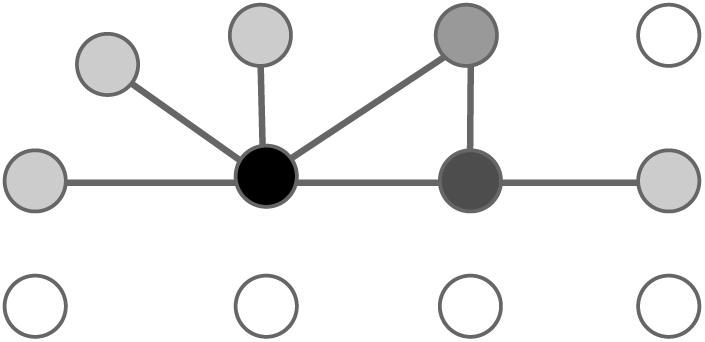
Network example where the darker the color of a node, the higher is its degree.

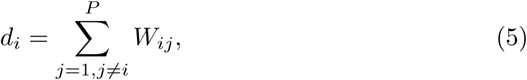

where *W*_*ij*_ are the elements of a positively weighted adjacency matrix.

Nodes with high degree are called *hubs*, while the nodes with small degree, either disconnected or with few edges, will here be denoted as *orphans*. These types of nodes play different roles in the network, as hubs can be used to reduce the diameter of a network and have a greater influence over its highly populated neighborhood. On the other hand, orphans are more isolated and play a smaller role in the overall or local topology.

The Pearson’s correlation and covariance were used to build two different networks that are fully connected and then trimmed using a cut-off value that keeps only the edges with highest values. This threshold was experimentally adjusted from a set of 20 different cut-off values and tested using 10-fold cross validation. The results, however, were inferior to using independent information described below.

In addition to these data-dependent methods, we used the String network for the human genome, which records functional protein-protein interactions that have been observed experimentally and estimated computationally (Szk-larczyk et al. 2017). The String network was downloaded from the project’s web page^2^, and filtered to use the *Homo sapiens* interactions only and to exclude text-based scores. Each edge of the networks denotes the overall score between the pair of proteins. As each protein can be mapped to one or more genes, we used the Ensembl genome database (Ensembl 2017) to determine this mapping and produce a gene network.

## 3 Contributions

The DegreeCox method had several shortcomings as it only implemented the *L*2-norm in the regularization function. This allows to converge to a solution with a small norm, however, the trained models then select every feature available. Moreover, the dual problem was used to solve the optimization problem in model inference, leading to an approximation of the maximum partial log-likelihood. We now propose a new framework using the elastic net approach that includes both the *L*1 and *L*2-norms, thus making this method sparse. Additionally, we are testing the reverse hypothesis that promotes nodes that have few or no edges in the network, thus introducing OrphanCox.

The degree of the nodes was calculated using a preprocessed network that eliminated edges below a given cut-off value. The resulting degree vector was then scaled between 0 and 1, and transformed using a double exponential heuristic function to increase the difference between the nodes with small degree and those with high degree:

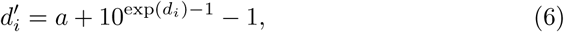

where *a* prevents disconnected nodes to have *weight* = 0 and *d*_*i*_ is the degree of node *i*. This expression can be further generalized from the equation below that has some similarities with the properties of the Generalized Extreme Value distribution (Gumbel 1935):

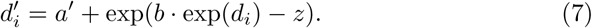

This transformation is an experimental requirement, as simply using the original degree showed little difference in the selected features between a classical elastic net run and network-based one. As such, this transformation preserves the ranking of the degree while expanding the difference as the degree increases.

### 3.1 HubCox

We propose HubCox as an extension to DegreeCox that includes the *L*1-norm that will allow the model inference to not only find the coefficients that minimize Equation 4 but also performs feature selection (Tibshirani 1996). This method takes advantage of existing implementations (Friedman, Hastie, and Tibshirani 2010) that use the coordinate descent algorithm to solve this optimization problem.

The regularization function **Ψ** is described in Equation 4. It penalizes nodes with small degree (Fig. reffig:sparse) by using as *w* the following expression:

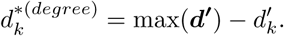

### 3.2 OrphanCox

We also introduce OrphanCox as a method that promotes nodes that are disconnected or with few edges in a network (Fig. 2). This is done by penalizing hubs in the regularization function **Ψ** with a higher weight. The hypothesis behind OrphanCox is that hubs have a regulatory effect and a high correlation with adjacent nodes and, as such, the variability in the data between the hub and less connected nodes could explain the response, *i.e.*:

**Figure 2:**
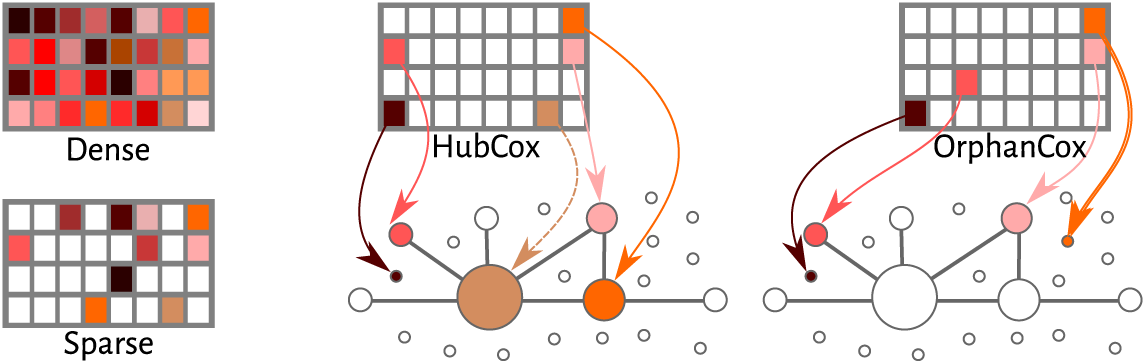
Models representing dense solutions without regularization, sparse, promoting hubs and promoting orphans.

**Figure 3:**
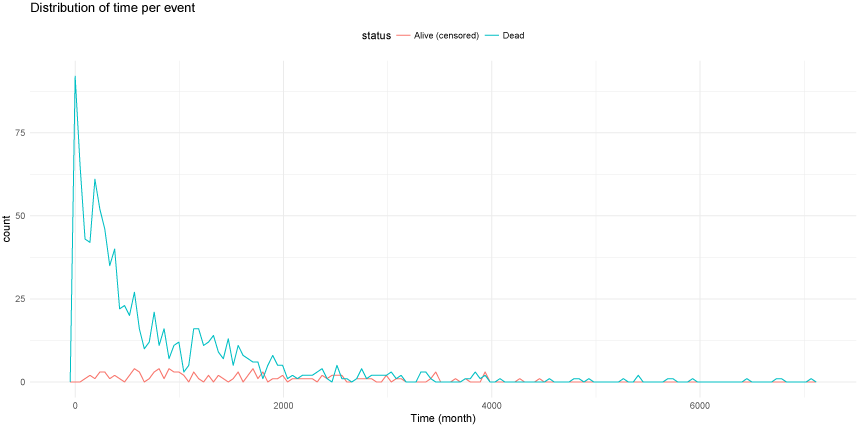
Event frequency distribution over follow-up time.

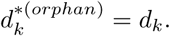

### 3.3 glmSparseNet **R-package**

An R-package called `glmSparseNet` was developed to include the methods described in this paper. This package depends on the coordinate descent implementation of the `glmnet` R-package (Friedman, Hastie, and Tibshirani 2010) to minimize the negative partial log-likelihood problem for the survival models and allows to define the weights calculated for both HubCox and OrphanCox. It also implements efficient methods to calculate the correlation and covariance networks over large data sets by paralleling the work and having a low memory footprint.

This R-package implements not only Cox Proportional hazard models, but also every other distribution family supported by `glmnet`, namely *gaussian*, *poisson*, *binomial*, *multinomial*, *cox*, and *mgaussian*. It adds two new main functions called `glmSparseNet` and `cv.glmSparseNet` that extend both model inference and model selection via cross-validation with network-based regularization. These functions are very flexible and allow to transform the penalty weights after the centrality metric is calculated, thus enabling to change how the regularization is affected. To make things easier for the users, we made available a function, called `glmHub`, that penalizes weakly connected nodes in the network, and another called `glmOrphan` that penalizes hubs.

An additional R-package is available with the necessary scripts and functions to reproduce the results of the present work. It is available to download at github^3^ under license GNU GPL version 3. This support package requires a modified `glmnet` R-package that is up to three times faster at calculating cross-validation for the data sets used in this work. This modified package can be installed directly from its github repository^4^.

## 4 Dataset

The Breast Invasive Carcinoma (BRCA) data set from The Cancer Genome Atlas (TCGA) was used to test the methods described in this manuscript, in particular, we used gene expression levels and clinical data from patient follow-up consultations that indicated whether the patient was deceased, our event of interest.

The BRCA data comprise information from 1047 individuals and 55867 genes, from which 19868 are protein coding genes that are used in the analysis. The expression levels were already processed by TCGA using Fragments Per Kilobase Million (FPKM) expression units. The `brca.data` R package dis-tributes the clinical and expression levels for BRCA and is available for download at github^5^.

Before survival analysis, a preprocessing step was applied to the BRCA dataset, including: the exclusion of genes that did not have any variation in the expression levels and non-coding genes identified by the Ensembl genome database (Ensembl 2017) and the Consensus CDS (CCDS 2017) projects; the removal of individuals with missing follow-up time for censored observations and individuals with negative follow-up time for both censored and non-censored observations; the removal of duplicate samples per individual by keeping the first sample; the retention of samples uniquely from primary solid tumours; the normalization of the FPKM values log2 and Z-score transformations.

## 5 Results and Discussion

The methods described in section 3 were compared with a classical approach using elastic net and one of the largest cancer dataset available from TCGA, the BRCA. We used the String network in the penalization of HubCox and OrphanCox as the network is independent from the data set and avoids introducing bias in the models. These numeric experiments were performed by running a 10-fold cross validation fitting the whole data set using several *α* parameters and *(1)* Elastic net (EN); *(2)* HubCox; *(3)* OrphanCox. The value of 0.2 for *α* was chosen as it minimizes the two goodness-of-fit measures (Log-rank test and c-index) described below over the different parameters that were tested, and it provides a good dimensionality reduction without degrading the performance of the model.

Since different model selection strategies may lead to a distinct optimal number of selected features, we started by running a 10-fold cross-validation for each model to chose the best *λ* parameter for each one, by selecting the one with the minimum log-likelihood deviation. We then tuned the *λ* parameter to force similar model sizes in the other two methods in order to promote an unbiased comparison with similar model sizes. This results in having three *λ* parameters for each of EN, HubCox and OrphanCox, the optimal one calculated via cross-validation, herein called the base model, and two others that were tuned to match the same number of selected variables called complementary. Figure 4 shows the diagram of the workflow.

**Figure 4:**
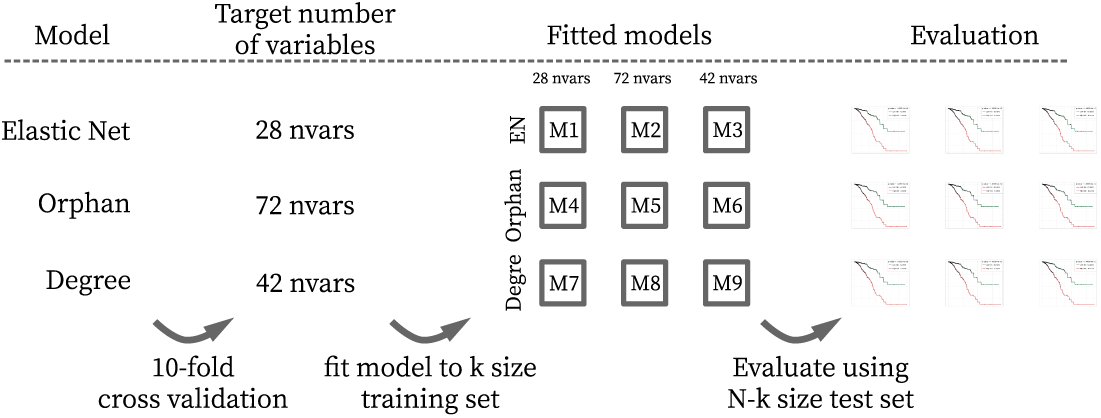
Diagram of benchmark framework in BRCA dataset.

The dataset was divided into a training and a test data set, comprising 80% and 20% of the data respectively. The results were assessed using two different criteria to evaluate the accuracy of the fitted relative risk of each individual. We used the concordance c-index (Harrell et al. 1982) that compares each pair of individuals from the population and checks if the fitted relative risks of the pair is concordant with the survival time, *i.e.* the individual with a lower relative risk survives longer than the other. The second evaluation metric is performed by separating the population in two groups by the median of the fitted relative risks. This allows to perform the Log-Rank test via the Kaplan-Meier estimator (Kaplan and Meier 1958), and to assess if the survival curves of the two groups are separable by comparing the p-values. Additionally, we also observed the survival rate of the long-term survival of the low risk group.

We start by looking at a single partition of the training and test data sets, fixed with a pseudo-random initial seed of 1985. Table 1 summarizes the results for the log-rank test and c-index metrics for the fitted models. We can observe from this single example that no method stands outs out of the three. While in the c-index metric, EN always is slightly higher then the other methods, with mixed results on the log-rank test.

**Table 1:** Results for a single partition of the test and training sets using String network. Lines with * have models trained with a target of the same number of variables as the base model.

| Base model to target $\lambda$ | Model | Selected Variables | Log-rank test | | C-Index | |
| --- | --- | --- | --- | --- | --- | --- |
|  |  |  | Train set | Test set | Train set | Test set |
| EN | EN | 84 | 0 | 0.0049797 | 0.8811822 | **0.6789366** |
|  | HubCox* | 83 | 0 | **0.0029993** | 0.8763837 | 0.6635992 |
|  | OrphanCox* | 85 | 0 | 0.0238635 | 0.87262124 | 0.6574642 |
| HubCox | EN* | 40 | 0 | **0.0277568** | 0.803588 | **0.6666667** |
|  | HubCox | 39 | 0 | 0.0732807 | 0.7916462 | 0.6451943 |
|  | OrphanCox* | 42 | 0 | 0.100175 | 0.8088773 | 0.6390593 |
| Orphan | EN* | 103 | 0 | 0.0120689 | 0.9051202 | **0.6768916** |
|  | HubCox* | 106 | 0 | 0.0080043 | 0.9022302 | 0.6595092 |
|  | OrphanCox | 105 | 0 | **0.0019162** | 0.8939419 | 0.6697342 |

Looking at the best models for each type in Figure 5, we can observe that the fitted survival curves of OrphanCox provide the best separation between low and high risk individuals and, very importantly, a much higher long term survival rate for the low risk individuals just short of 60%, while EN has around 40%. This statement still holds when looking at the EN model with 103 selected variables, which might indicate that this could be an improvement of network-based models over classical EN.

**Figure 5:**
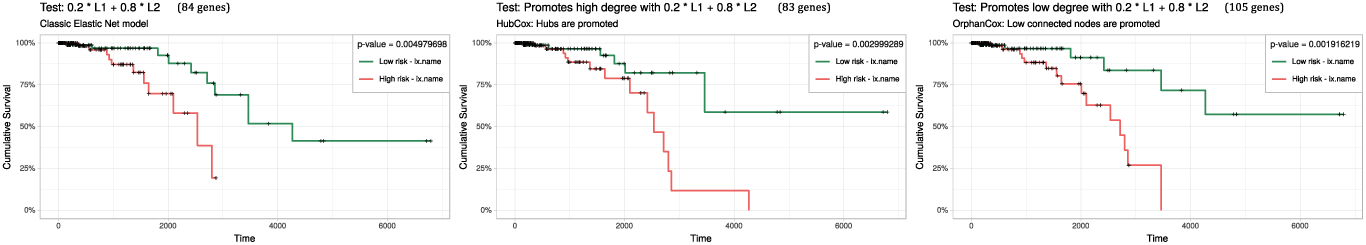
Survival curves for best models from all the models calculated with the same train and test partition with *α* = 0.2 and EN (left with 84 variables), HubCox (center with 84 variables) and OrphanCox (right with 105 variables) for low and risk groups separated by median relative risks.

When studying the stability of selected genes over 1000 resampling of the train and test sets, we observe that HubCox is the most consistent with 657 unique variables selected at least once over all the runs. OrphanCox and EN perform much worst with 1129 and 1339, respectively. Figure 6 shows the Venn diagram of the overlapping genes between models. These results are for the models fitted using the EN base model, with a target of selected genes of 83.

**Figure 6:**
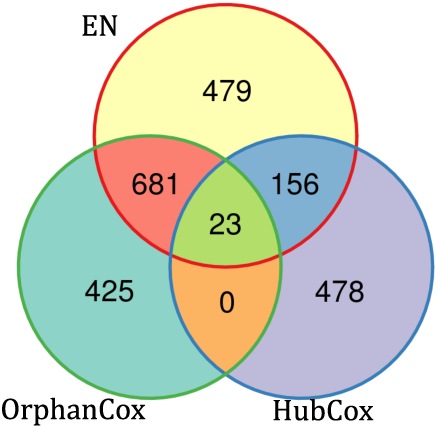
Venn diagram of the overlapping selected genes of all 1000 fitted models. Showing diagram for the EN base model that has a target of 83 selected genes.

It is interesting to note that there is a bigger overlap of selected genes between OrphanCox and EN, while the genes found by HubCox almost do not intercept. This latter pattern is something to be expected as the underlying hypothesis for each of the network-based model is inverse. Nevertheless, there are 23 genes that were commonly selected, such as *OR5AU1* (protein part of the olfactory receptor proteins) and *CEL* (a glycoprotein secreted from the lactating mammary gland into human milk), that have degrees of 325 and 251, respectively (network where the highest degree is 10394 and median degree value is 220). Despite these genes being heavily penalized in HubCox, they are still selected as their inclusion outweighs the network penalty.

By further looking into the genes selected, we used the model with the same *λ*, but now trained with the full BRCA dataset to understand which genes are selected by the network-based methods and not by EN. In part of this analysis we used the Cancer Hallmarks Analytics Tool (Hanahan and Weinberg 2011), denoted as CHAT, to query the selected genes against a database of known functions related to cancer. The CHAT database was populated based on a text-mining analysis of 26 million PubMed abstracts. Figure 7 shows a heatmap of the hallmarks counts per gene, per model, with some genes appearing few times (in light blue) and some highlighted (in dark blue). Looking at the highlighted ones, there are some very well known genes related to cancer, such as *TP53* (Walsh et al. 2006), *NRAS* (Cimino et al. 2012) and *KRAS* (Paranjape et al. 2011). However, all of these were selected by HubCox, while OrphanCox selected genes that seemed less studied and discussed. Ultimately, this is directly associated with the network and the hypothesis being tested with the regularizer, as known genes are bounded to have a more central role in the network, especially if they have been studied as associated with multiple cancer types. In particular, we observe in the results that HubCox promotes these genes in the solution, making this method more suitable to have a stable selection of variables and an improved performance. In contrast, OrhanCox promotes unknown or little studied genes that can become good candidates to further research. To conclude, these two comparably performing models might be seen as complementary and suited to different applications.

**Figure 7:**
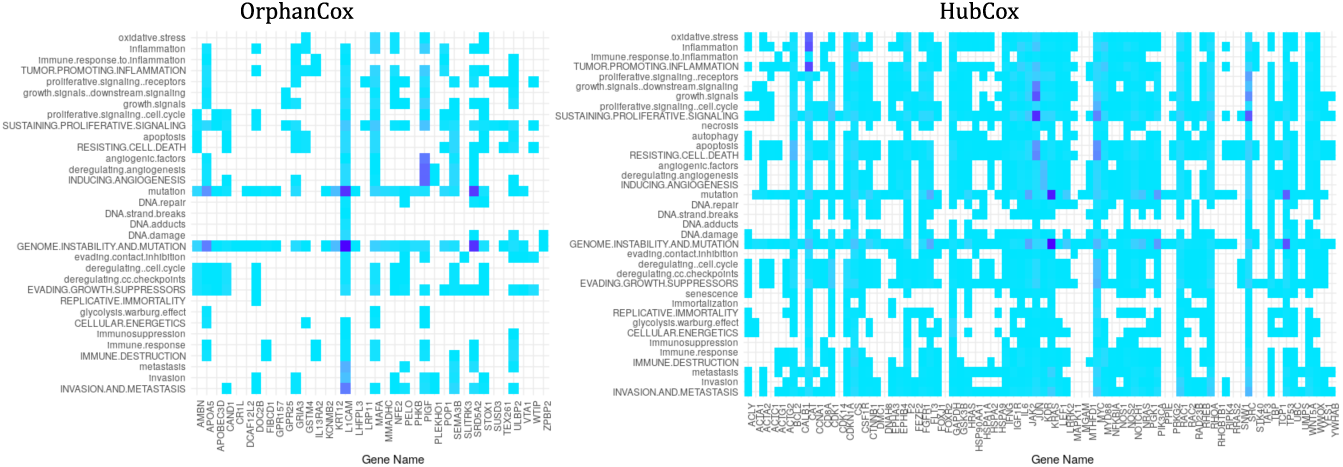
Counts of hallmarks association in CHAT database, the darker the cell, the higher the count of hallmarks for a particular gene. Only showing genes that do not intercept with EN.

## 6 Conclusion

The initial hypothesis of this work was that the enrichment of survival and generalized linear models with *a priori* network information would improve the models’ predictive quality, stability and interpretability. Our results show that the models’ goodness-of-fit metrics is comparable with elastic net, and seems to have a better performance in predicting long term survival individuals. Moreover, we have achieved a good stability of selected variables in resampling tests, which is crucial for models’ interpretability. Both the promotion of hubs or low connected nodes have their merits, and while it would be interesting to have a definitive model for transcriptomic breast cancer data, the results obtained from both methods are suitable for different purposes.

The variability present in transcriptomic data allows for models with different combinations of genes to be equivalent. These proposed methods enable a more guided approach for gene selection and biomarker identification in precision medicine.

## 7 Acknowledgments

The authors thank Niko Beerenwinkel and Wolfgang Huber for fruitful discussions. This work was partially supported by the European Union Horizon 2020 research and innovation program under grant agreement No. 633974 (SOUND project), and by national funds through Fundaçã,para a Cinêcia e a Tecnologia (FCT) with reference UID/CEC/50021/2013 (INESC-ID), through IDMEC, under LAETA, project UID/EMS/50022/2013, and PERSEIDS (PTDC/EMS-SIS/0642/2014), SFRH/BD/97415/2013 and IF/00653/2012.

## Footnotes

1 https://github.com/sysbiomed/glmSparseNet

2 https://string-db.org/cgi/download.pl?sessionId=VPkN7How44B1&species_text=Homo+sapiens

3 https://github.com/averissimo/glmSparseNetPaper

4 https://github.com/averissimo/glmnet

5 https://github.com/averissimo/tcga.data/releases/download/2016.12.15-brca\/brca.data_1.0.tar.gz

